# Automated Adherent Cell Elimination by a High-Speed Laser Mediated by a Light-Responsive Polymer

**DOI:** 10.1101/280487

**Authors:** Yohei Hayashi, Junichi Matsumoto, Shohei Kumagai, Kana Morishita, Long Xiang, Yohei Kobori, Seiji Hori, Masami Suzuki, Toshiyuki Kanamori, Kazuhiro Hotta, Kimio Sumaru

## Abstract

Conventional cell handling and sorting methods require manual dissociation, which decreases cell quality and quantity and is not well suited for monitoring applications. To purify adherent cultured cells, *in situ* cell purification technologies that are high throughput and can be utilized in an on-demand manner are expected. Previous demonstrations using direct laser-mediated cell elimination revealed only limited success in terms of their usability and throughput. Here, we developed a Laser-Induced, Light-responsive-polymer-Activated, Cell Killing (LILACK) system that enables high-speed and on-demand adherent cell sectioning and purification. This system employs a visible laser beam, which does not kill cells directly, but induces local heat production through the trans-cis-trans photo-isomerization of azobenzene moieties in only the irradiated area of a light-responsive thin layer. Using this system in each passage for sectioning, human induced pluripotent stem cells (hiPSCs) were maintained their pluripotency and self-renewal during long-term culture. Furthermore, combined with deep machine-learning analysis on fluorescent and phase contrast images, a label-free and automatic cell processing system was developed by eliminating unwanted spontaneously differentiated cells in undifferentiated hiPSC culture conditions.

## Introduction

The purification of different types of cultured cells is critical in various biomedical fields, including basic research, drug development, and cell therapy. Conventionally, fluorescence-activated cell sorting (FACS), affinity beads (e.g., magnetic-activated cell sorting (MACS)), gradient centrifugation, and elutriation have been used for cell purification^1^. However, these technologies are essentially aimed at floating cells in suspension. For adherent cells, the process of detaching, dissociating, sorting, and reseeding can result in low yield and in altered cell characteristics^2^. For adherent cells, antibiotics or special chemicals have limited use for the selection of genetically modified cells or specialized nutrient-requiring cells, respectively.

To purify adherent cultured cells, *in situ* cell purification technologies that are high throughput and can be utilized in an on-demand manner are expected. Since light irradiation can be precisely controlled by computers on a microscopic scale and is suitable for sterile processes, methodologies using light have been examined to automate this operation. Among these methods, laser-mediated cell elimination is a promising technology^3^. Previous demonstrations, however, revealed only limited success of this method since it requires a high amount of energy to eliminate or move the cells directly, resulting in moderate speed of processing (∼1,000 cells/sec)^3^-^6^. This high auxiliary energy input produces an enormous amount of heat that kills surrounding cells, which destroys the focusing of cell processing. Also, the heat might denature the components in culture media. We previously demonstrated that killing cells through the microprojection of visible light by using photo-acid-generating substrates^7^,^8^. However, one projection covered only 0.1 cm^2^, and the cell elimination took longer than 1 minute in these previous studies.

To overcome these limitations, we have developed a Laser-Induced, Light-responsive-polymer-Activated, Cell Killing (LILACK) system enabling high-speed and on-demand adherent cell sectioning and purification (schemes shown in Figure 1a). This LILACK system employs a visible laser beam with a 405 nm wavelength, which does not kill cells directly, but induces local heat production in only the irradiated area of a light-responsive thin layer composed of poly[(methyl methacrylate)-co-(Disperse Yellow 7 methacrylate)]. The energy of the irradiated laser is converted to heat efficiently through the trans-cis-trans photo-isomerization of azobenzene moieties, without photolysis of the polymer^9^. Further, the polymer is free from fluorescence emission and absorbance in most of the visible range, which hinders cell observations. Using this system, human induced pluripotent stem cells (hiPSCs)^10^,^11^ were sectioned in each passage to maintain their pluripotency and self-renewal in long-term culture. Furthermore, combined with deep machine-learning analysis on fluorescent and phase contrast images, a label-free and automatic cell processing system was developed by eliminating unwanted spontaneously differentiated cells in undifferentiated hiPSC culture conditions. This LILACK system enables to select adherent cells *in situ* on an acceptable timescale using the precise and very fast scanning of a well-focused visible laser through a light-responsive polymer layer, and automatic label-free cell purification combined with efficient imaging analysis based on deep machine learning methods.

**Figure 1.**
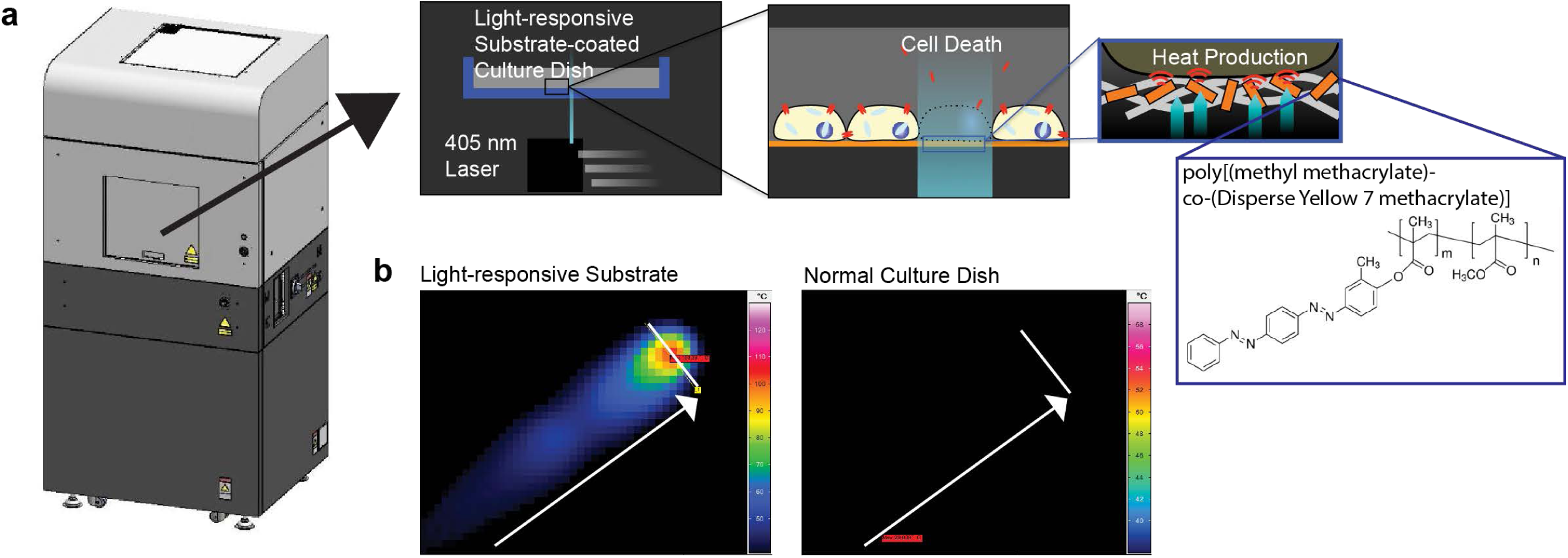
Schemes of the LILACK system and its focused heat production. (a) Schemes of LILACK system. (b) Thermal images of the surfaces of cell culture dishes after laser irradiation. The laser was irradiated at 80 mm/sec and 0.3 W with a width of 50 µm towards the arrow direction. Left: light-responsive polymer-coated dish. Right: normal cell culture dish. The bar in the thermally responsive area indicates 50 µm.

## Results

### Focused heat production by LILACK system

First, we examined the effectiveness of local heat production through the trans-cis-trans photo-isomerization of azobenzene moieties. Laser irradiation at 0.3 W and 80 mm/sec and with a diameter of 50 µm generated heat at more than 50 °C over focused area of the light-responsive-polymer-coated dishes accurately. In contrast, laser irradiation with the same conditions did not generate detectable heat on the surface of normal cell culture treated dishes (Figure 1b and S1). These results indicated that this scheme enables effective cell killing even at very fast beam scanning without damaging neighbouring unirradiated cells.

### Growth and viability of hiPSCs on the light-responsive polymer

We examined growth and viability of hiPSC cultured on the light-responsive polymer in both on-feeder^10^ and feeder-free^12^,^13^ culture conditions. Growth and viability of hiPSCs on the light-responsive polymer were comparable to those on normal cell culture-treated substrates (Figure S2). These results indicated that the light-responsive polymer does not influence the cultured cell growth and viability.

### Cell killing effectiveness of LILACK system

We examined the induction of cell death by laser scanning at different power settings and 100 mm/s with a diameter of 50 µm. Laser scanning at a power of 0.8 W or higher readily induced hiPSC death over a diameter of approximately 50 µm (Figure 2a). Also, using a 4.4 W laser, a scanning speed of 1,000 mm/s or higher readily induced hiPSC death. We obtained similar results with the Madin-Darby canine kidney (MDCK) cell line^14^ (Figure 2b). In the laser scanning at different power setting and 100 mm/s with a diameter of 50 µm, a power of 0.8 W or higher readily induced MDCK cell death over a diameter of approximately 50 µm. Also, the combinations, 2.5 W with 500 mm/s, 2.0 W with 400 mm/s, 1.6 W with 320 mm/s, and 1.0 W with 200 mm/s, which should give the same energy intensity, readily induced similar patterns of MDCK cell death. Using 5.0 W laser, a scanning speed of 1,000 mm/s or higher readily induced MDCK death. Assuming that one cell has 10 µm in diameter, we have demonstrated that this LILACK system enables to process >10,000 cells/s in throughput.Furthermore, over 100,000 cells/s can be processed by the laser irradiation with the higher power laser. These results demonstrate that our LILACK system enables high-speed and on-demand laser-mediated cell death of various adherent cultured cells.

**Figure 2.**
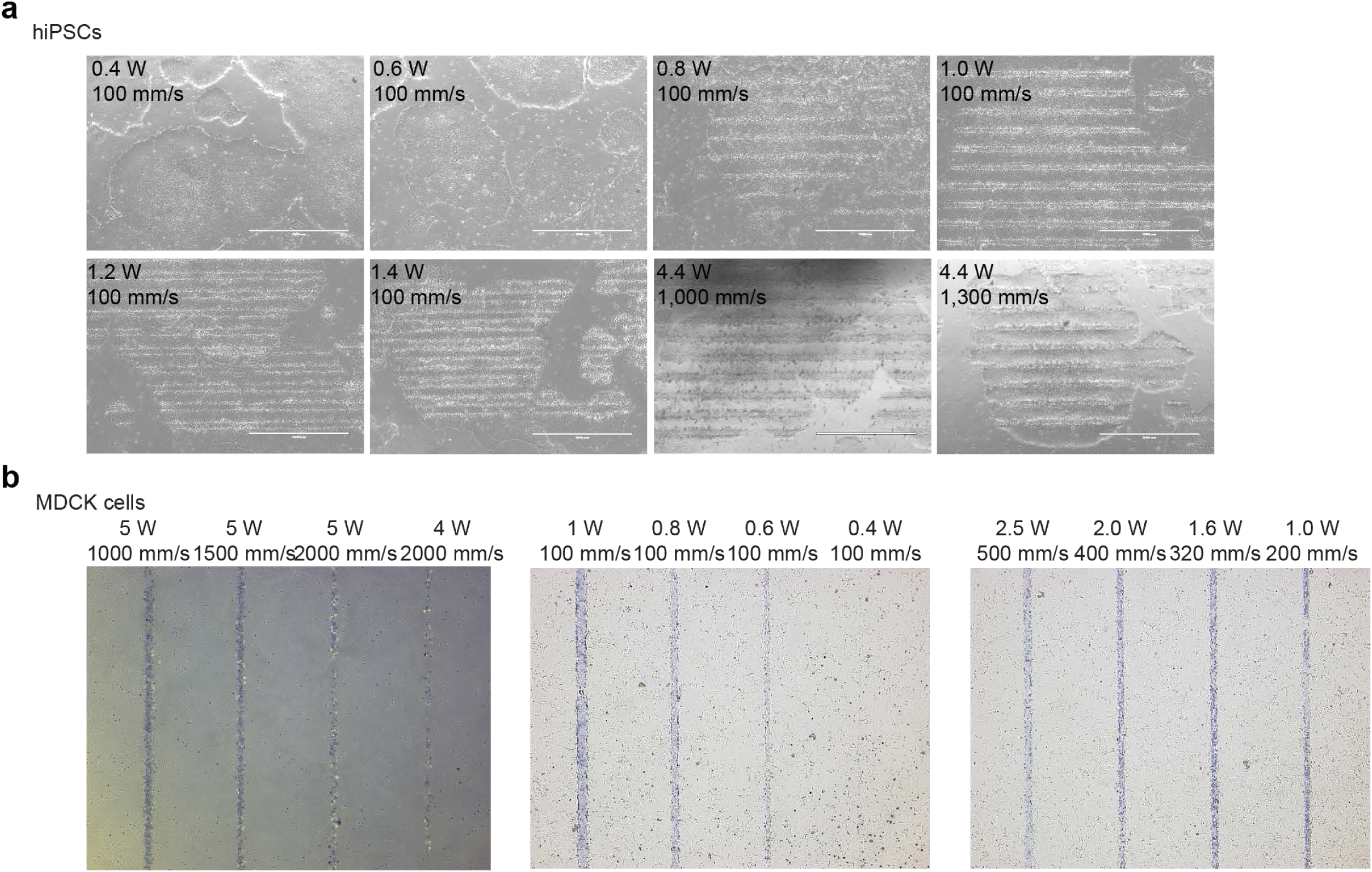
Cell killing of LILACK system on hiPSCs and MDCK cells. (a) Phase-contrast images of hiPSCs after scanning by lasers with different power and speed settings. Scale bars are 1.0 mm. (b) Phase-contrast images of MDCK cells treated with 0.4% trypan blue solution after laser scanning at different power and speed settings. Dead cells were positively stained with trypan blue. The intervals between lines were 1 mm.

### Long-term hiPSC culture using LILACK system

To examine the feasibility of this LILACK system in regenerative cell therapies, we examined the effect of the long-term culturing of hiPSCs using LILACK system (demonstrated in Movie S1). hiPSC colonies cultured on the light-responsive polymer and on feeder cells were cut in a grid pattern by laser scanning and treated with cell dissociation enzyme to generate floating cell clumps (Figure 3a). The laser beam of 50 µm with 1.0 W intensity was scanned at the velocity 100 mm/s, with 450 µm intervals. Although extra time was needed for the interval between linear scans, the grid scanning took only 86 sec to section the cell monolayer in whole area of the 35 mm culture dish surface (9 cm^2^). These hiPSC clumps were transferred to a new light-responsive-polymer-coated culture dish with feeder cells, and the culture continued. After 10 passages through the system consisting of the above steps, the hiPSCs were characterized. The karyotype of these hiPSCs was maintained in all the cells (i.e., 50 cells/50 cells) (Figure 3b). These hiPSCs expressed self-renewal markers of pluripotent stem cells, such as NANOG, OCT3/4, SSEA4, and TRA1-60, which were determined immunocytochemically (Figure 3c). When differentiated into three germ layers using embryoid body (EB) formation, these cells expressed, microtubule-associated protein 2 (MAP2) as an ectoderm marker, smooth muscle actin (SMA) as a mesoderm marker, and α-fetoprotein (AFP) as an endoderm marker, detected by immunocytochemistry (Figure 3d). Additionally, these hiPSCs were free of harmful viruses and mycoplasma detected by quantitative PCR-based tests (Table S1). These results indicate that our LILACK system using laser-mediated cell processing maintained the self-renewal, pluripotency, and biological safety of hiPSCs during long-term culture.

**Figure 3.**
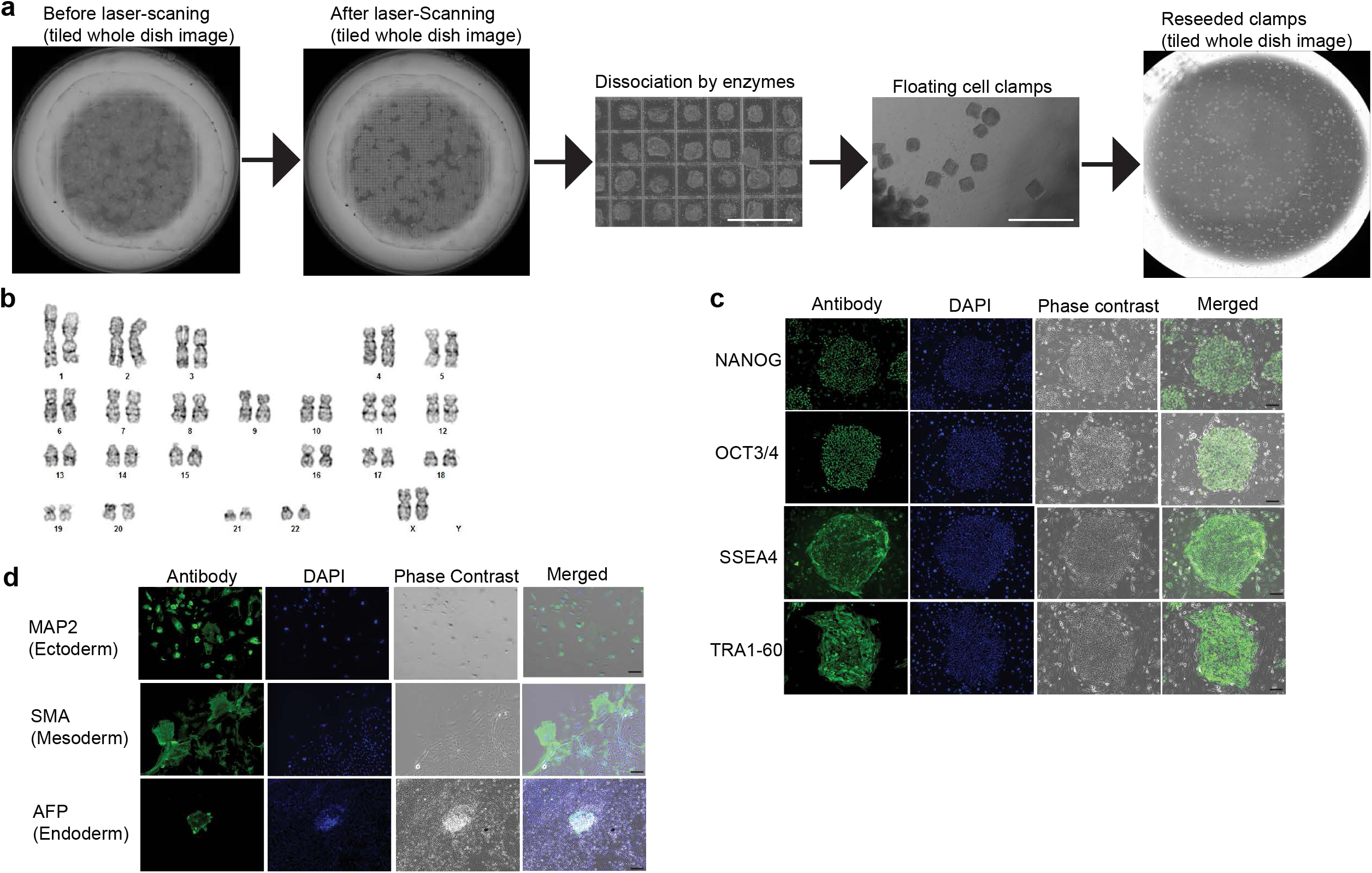
Long-term culture of hiPSCs using LILACK system. (a) Scheme of the sectioning of a hiPSC monolayer by grid-patterned laser scanning and subcultures. Scale bars, 1 mm. (b) Karyotype example of hiPSCs that underwent 10 passages through the system after laser-mediated sectioning. (c) Self-renewal marker expression in hiPSCs that underwent 10 passages through the system after laser-mediated sectioning. Scale bars, 100 µm. (d) Expression of three germ layer markers in the EBs differentiated from hiPSCs that underwent 10 passages through the system after laser-mediated sectioning. Scale bars, 100µm.

### Deep learning analysis to identify spontaneously-differentiated hiPSCs

Because this LILACK system enables cell processing based on phase-contrast and fluorescent cell morphologies, we developed a label-free cell elimination system based on deep machine-learning imaging analysis. Emerging spontaneous differentiated cells in undifferentiated human pluripotent stem cell culture conditions is important problems in obtaining reproducible and reliable results in basic scientific studies and therapeutic cell production^15^-^17^. We tried to solve this problem by automatically eliminating these differentiated cells using this LILACK system combined with deep learning imaging analysis. First, we collected phase-contrast and fluorescence images using a fluorescent probe, rBC2LCN-FITC, to mark undifferentiated hiPSCs^18^,^19^ (example in Figure 4a). Using the LILACK device, which carries the function to collect tiled images of a whole dish, 12,556 differentiated images and 18,834 undifferentiated images were collected. These images were used to train a convolutional neural network (CNN). Figure 4b shows the structure of the CNN. The size of an input image was set to 70×70 pixels. Since the size of an original image is 1920×1200 pixels, local regions of 70×70 pixels can be cropped without overlap, and the regions can be fed into the CNN. Our CNN consisted of two convolutional layers, two pooling layers, and a fully connected layer. Thirty-two filters with 5×5 and 3×3 kernel sizes were used in the first and second convolutional layers, respectively. After the convolutional layers, we used a maximum pooling with 3×3 and 2×2 kernel sizes. The softmax cross entropy loss was used to train the CNN. Undifferentiated images were gathered in two ways. First, 70×70 local regions were cropped randomly from differentiated and undifferentiated regions, and the CNN was trained by those images. The trained CNN was applied to images, and misclassified undifferentiated regions were gathered. The gathered undifferentiated cell images were added to the training images, and the CNN was trained again and used as the classifier. Figure 4c shows the original image and the detection result from the CNN. The probability of obtaining a differentiated class from the CNN is assigned to the pixel in the resulting image. The black pixel in the resulting image shows the differentiated region. The intensity of the black colour reflects the probability: darker colours indicate higher probability. These results indicated that deep machine-learning imaging methods were able to distinguish spontaneously differentiated cells in undifferentiated hiPSC culture conditions.

**Figure 4.**
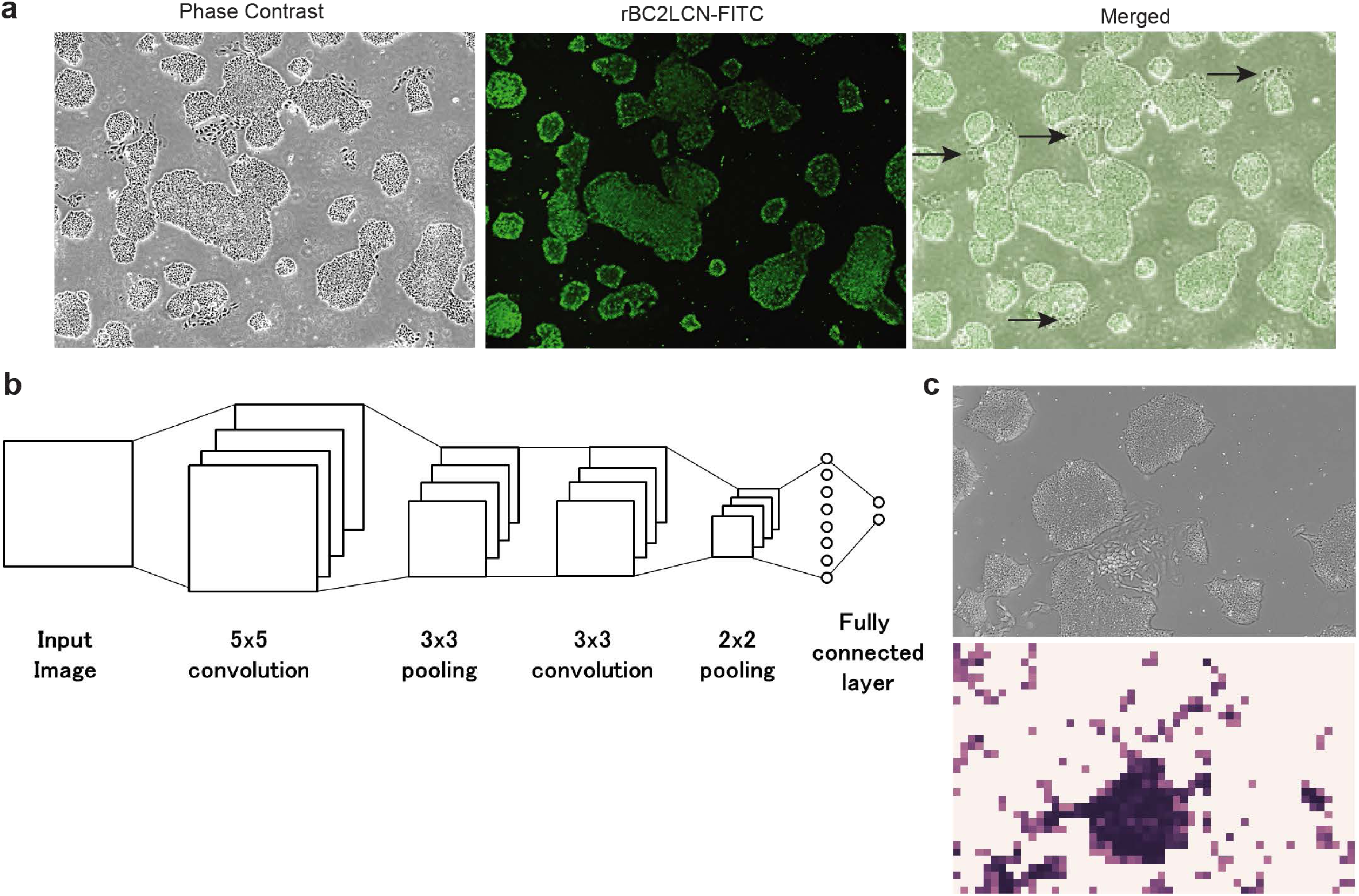
Deep machine learning to identify spontaneously-differentiated hiPSCs. (a) Example of the “teacher” image sets used in the machine learning. Live cell phase-contrast and fluorescence images from the undifferentiated marker probe (rBCN2CL-FITC) were combined. Arrows indicate the spontaneously differentiated cells that were rBCN2CL-negative. (b) Scheme of the CNN structure. (c) Example of an input image (upper) and output image (bottom) generated by the algorithm generated from the CNN results. In the output image, dark areas indicate differentiated cells automatically classified from the input image.

### Automatic in situ purification of undifferentiated hiPSCs using LILACK system

Last, we applied this trained classification algorithm to our laser-mediated cell elimination with only phase-contrast images (Figure 5a). The laser beam of 50 µm with 0.5 W intensity was scanned at the velocity 100 mm/sec, with 25 µm intervals. The laser irradiation was activated only when the beam was in the target areas. Purification in whole area of the culture surface took 667 sec, which could be shorten according to the distribution of the target cells. hiPSCs, which were treated with automated cell elimination or not, were collected and analysed for TRA1-60-positive cells, which indicated undifferentiated hiPSCs^20^,^21^, by flow cytometry analysis. TRA1-60-positive cell ratio increased to 97% or more after laser irradiation to eliminate the “differentiated” cells classified by this algorithm (Figure 5b and S3). These results indicate that *in situ* label-free cell purification was automatically achieved using our LILACK system combined with imaging analysis based on deep learning.

**Figure 5.**
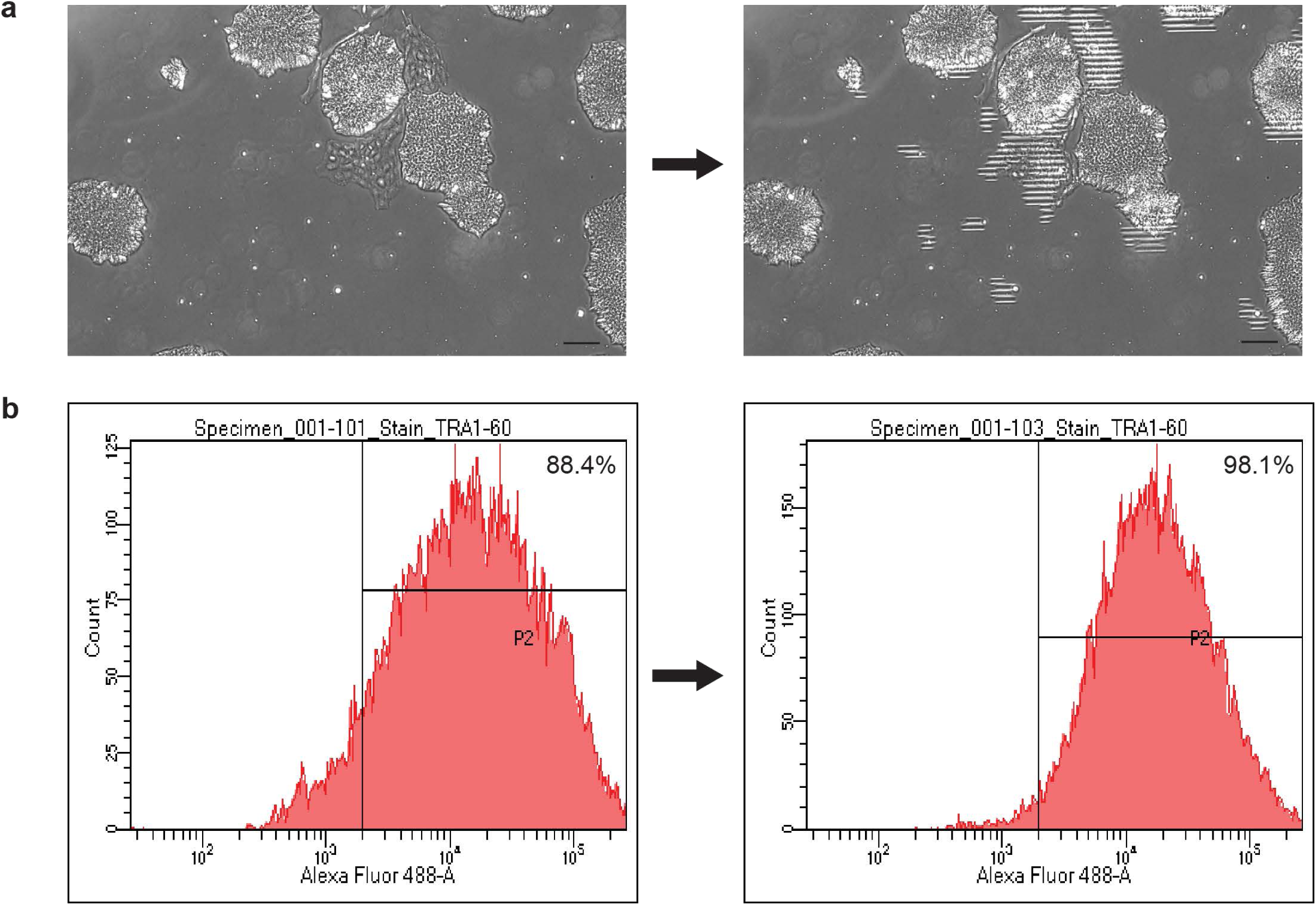
*In situ a*utomatic purification of undifferentiated hiPSCs using LILACK system (a) Example of the automatic laser-mediated elimination of spontaneously differentiated cells in normal hiPSC culture conditions. Scale bars, 100 µm. (b) Flow cytometric analysis of TRA1-60, a cell surface pluripotency marker, expression in hiPSCs with or without laser-mediated cell elimination of spontaneously differentiated cells.

## Discussion

We have developed a novel system for selecting adherent cells *in situ* on an acceptable timescale using the precise and very fast scanning of a well-focused visible laser through a light-responsive polymer layer at 100 mm/sec or more (>10,000 cells/sec). Combined with efficient imaging acquisition and analysis based on machine learning, we achieved automatic label-free cell purification and sectioning. Compared with conventional methods, such as FACS, affinity beads, gradient centrifugation, elutriation, and direct laser-induced cell killing (summarized in Table 1), our LILACK system may have advantages in maintaining the cell numbers and characteristics since our methods do not directly affect the un-irradiated cell activities. Thus, we believe that our methods can be widely used in various biomedical fields, including basic research, drug development, and cell therapy.

**Table 1.** Summary of the comparisons between conventional cell purification systems and our new LILACK system.

| **Selection Type** | **FACS** | **Affinity Beads** | **Gradient Centrifugation** | **Antibiotics-based cell separation** | **Direct Laser Cell Killing System** | **(Indirect) LILACK system** |
| --- | --- | --- | --- | --- | --- | --- |
| **Cell Preparation** | Dissociation | Dissociation | Dissociation | Genetic modification with antibiotics-resistant genes or cell type specific chemicals | No need (or labeled by fluorescent probes) | No need (or labeled by fluorescent probes) |
| **Culture condition needed** | None | None | None | Specific antibiotics or chemicals | Specific dye and only glass dishes | Specific light-responsive-substrate-coated dishes |
| **Throughput** | ~1,000 cells/sec | ~ten million cells/sec | ~ 30 minutes/centrifuge | Several days or weeks | ~1,000 cells/sec | >10,000 cells/sec |
| **Device** | FACS (BD, Beckman Coulter and others) | Magnetic Sorting Device (Miltenyl Biotec and others) | Centrifuge and gradient reagents (many manufacturers) | None | LEAP (Intrexon) and others | Specific Laser-scanning device (Kataoka Corp) |

The LILACK system should be particularly suitable for use in cell therapies in future. Because our LILACK system does not require direct manual handling to process cells and can be easily operated in an isolated space or even in an incubator for cultured cells in principle, it can reduce risk of contamination or exposure to hazardous substances compared with the risks associated with conventional cell sorting using FACS, MACS, or other filtering methods. Additionally, compared with previous reports using specific dyes supplemented in the cell culture medium for the efficient energy absorption of laser irradiation^3^-^6^, our methods do not produce free chemicals in the culture medium. Although the light-responsive polymers used in this method should be thoroughly scrutinized for biological safety, the LILACK system has great potential for use in cell therapies, with the capability for real-time monitoring and *in situ* automated cell processing combined with imaging analysis based on machine learning.

To achieve label-free cell purification of undifferentiated hiPSCs based on the colony morphology observed in phase contrast images, we utilized a deep-learning-based classification of undifferentiated and differentiated hiPSCs. Previous studies demonstrated the utility of morphological analysis to monitor the quality of hiPSCs^22^,^23^. In recent years, the effectiveness of deep machine learning for image recognition has been demonstrated. CNNs have worked well in various applications, such as object categorization^24^, object tracking^25^, action recognition^26^ and particle detection in intracellular images^27^. Therefore, we used a CNN to classify differentiated cells and undifferentiated cells in an unbiased manner. We successfully demonstrated that undifferentiated hiPSCs were enriched by the automated cell killing system based on the CNN-predicted programme to identify spontaneously differentiated cells. This demonstration indicates that the hiPSC quality could be continuously refined from non-labeled images combined with our laser-processing technology. Since our technology can be applied to any adherent cells and does not usually interfere with the fluorescence from labeled proteins used for various biological purposes, we believe that our technology is compatible with the purification of various fluorescent reporter cells, as well as non-labeled cells. Since the classification of cell types based on deep-learning methods is advancing rapidly, the importance and functionality of our technology will be further enhanced in the near future.

## Methods

### Fabrication of Photo-Responsive Culture Substrates

The photoresponsive polymer used in this study was poly[(methyl methacrylate)-co-(Disperse Yellow 7 methacrylate)] (#579149, Sigma-Aldrich), which was composed of ∼25 mol% azobenzene moieties. The polymer was dissolved in a mixture of 2,2,2-trifluoroethanol and 1,1,1,3,3,3-hexafluoro-2-propanol (10 wt%) to form a 1.0 wt% solution. Next, 20 µL of the solution was spin-coated on a cell culture dish with a 35 mm diameter (#3000-035, AGC Techno Glass, Japan) at 2,000 rpm under an N_2_ atmosphere, and then the substrates were annealed for 2 hours at 80 °C. The absorbance of the substrate was typically 0.25 at 405 nm and less than 0.05 at wavelengths greater than 500 nm.

### Thermal Imaging

Thermal images were taken by an infrared thermography camera, InfReC H9000 (Nippon Avionics, Japan) with a 5 µm microscopy lens with the following measurement parameters: wavelength, 2 ∼ 5.7 µm; temperature, −10 °C ∼ 1,200 °C; temperature resolution, 0.025 °C at 30 °C; frame rate, 200 Hz; spatial resolution, 5 µm; and temperature accuracy, 2% at an environmental temperature of 10 °C ∼ 40°C. The laser irradiated a cell culture dish coated with or without the light-responsive substrate at an environmental temperature of 29.5 °C without any liquid medium. The laser conditions were 0.3 W to 0.7 W at 80 mm/sec.

### Cell Processing by Laser Scanning

To achieve high-speed, automatic cell processing using 405 nm laser scanning, we developed an all-in-one apparatus that can take phase-contrast and fluorescence images and apply tiled, whole-dish laser-scanning in an on-demand manner (Kataoka Corp, Japan; Figure 1a). First, culture dishes are selected from a cell culture incubator, and photographs of the whole dish are taken and analysed. Then, the type (e.g., lattice pattern, rectangle, circle, and line) and conditions (e.g., speed and power) of laser scanning are selected. Once the conditions are determined, the on-demand laser irradiation is initiated. The irradiated culture dishes are then re-incubated and subjected to assays. A demonstration of the procedure is shown in Movie S1.

### MDCK Cell Culture

An MDCK cell line was purchased from the RIKEN BioResource Center. The cells were maintained in Eagle’s minimum essential medium (Sigma-Aldrich) supplemented with 10% fetal bovine serum (Gibco BRL). For passage through the system, the culture medium was removed and discarded. The cell layer was rinsed twice with a 0.25% (w/v) trypsin-0.53 mM EDTA solution to remove all traces of serum that contained trypsin inhibitor. The trypsin-EDTA solution was added to the dish, and the cells were observed under an inverted microscope until the cell layer dispersed (usually within 5 to 15 minutes). Complete growth medium was added, and the cells were aspirated by gently pipetting. Appropriate aliquots of the cell suspension were added to new light-responsive-polymer-coated dishes. The cells were incubated at 37°C and 5% CO_2_ in an incubator. The medium was replaced every 2 to 3 days. Laser irradiation experiments were performed on fully confluent conditions.

### hiPSC Culture

hiPSCs (201B7 or 1231A3 lines)^10^ were obtained from CiRA (Center of iPS Cell Research and Application) at Kyoto University through the RIKEN BioResource Center (Tsukuba, Japan). All media and reagents were purchased from commercial sources. For on-feeder culture conditions, the cells were maintained in Primate ES cell medium (Reprocell, Tokyo, Japan) or Stemsure on-feeder hPSC medium (Wako, Osaka, Japan) supplemented with 4 ng/ml bFGF and penicillin/streptomycin (Nacalai Tesque) on SNL feeder cells, which were from a mitomycin C (Sigma Aldrich)-treated SNL 76/7 cell line (Cell Biolabs) described previously^28^-^32^. For subculturing, the cells were detached from the culture dish using CTK solution (Reprocell)^33^. The SNL cells were cultured in fibroblast medium (DMEM supplemented with 10% fetal calf serum, 1% penicillin, and 1% streptomycin) on gelatin-coated dishes. These cell clumps were transferred to a new light-responsive-polymer coated dishes on SNL feeder cells at 1:3 to 1:10. For feeder-free culture conditions, the cells were maintained in StemFit AK02N medium (Ajinomoto, Tokyo, Japan) supplemented with all the abovementioned supplements on 0.5 µg/cm^2^ iMatrix511 (Nippi, Japan)-coated dishes^12^,^13^,^34^. A Rho-associated protein kinase (ROCK) inhibitor (Y-27632, Wako, Osaka, Japan) (10?µM) was added to the medium used in the passage process^35^. For subculturing, the hiPSCs adhered to a culture dish were washed with PBS and then treated with TrypLE Select (Thermo Fisher Scientific) in Dulbecco’s PBS at 37°C for approximately 5?minutes. After aspirating the solution, the cells were resuspended in fresh medium. Then, the cells were collected by cell scraping and pipetting. After counting the cell numbers, the cells were normally seeded at 2.5 x 10^3^ cells/cm^2^.

### Immunocytochemistry

For the immunocytochemical determination of pluripotency and differentiation markers, cells were fixed with PBS containing 4% (vol/vol) paraformaldehyde for 10 min at room temperature. The cells were permeabilized with PBS containing 0.1% Triton X-100 for 10 min at room temperature, then washed with PBS and treated with 1% BSA for blocking. For hiPSCs or reprogramming HDFs, the primary antibodies used were SSEA4 (0.5 µg/mL; eBiosciences), TRA-1-60 (0.5 µg/mL; eBiosciences), NANOG (2 µg/mL, AF1997; R&D Systems), OCT3/4 (1/200, sc-5279; Santa Cruz Biotechnology), MAP2 (1:1,000, AB5622; Millipore), a-SMA (1:1,000, A2547; Sigma), and AFP (2 µg/mL, MAB1368; R&D Systems).The secondary antibodies used were Alexa Fluor 488- or 555-conjugated goat anti-mouse IgG (1:200; Invitrogen), Alexa 488- or 555-conjugated goat anti-rabbit IgG (1:200; Invitrogen), and Alexa 488- or 555-conjugated donkey anti-goat IgG (1:200; Invitrogen). Nuclei were stained with the 4’,6-diamidino-2-phenylindole (DAPI) contained in the VectaShield set (Vector Laboratories). The samples were analysed in randomly selected images with BZ-X710 (Keyence, Osaka, Japan).

### HiPSC Differentiation

EBs were generated by CTK medium treatment of day-7 hiPSC cultures to remove colonies from the culture dishes. Colonies were grown in differentiation medium in a suspension culture using EZ-BindShut II dishes (Iwaki, Japan). hiPSCs were induced to spontaneously differentiate in a medium composed of StemSure DMEM (Wako) supplemented with 20% StemSure serum replacement (Wako), 2 mM L-alanyl-L-glutamine (Wako), 1% MEM non-essential amino acids (Wako), and 0.1 mM 2-mercaptoethanol (Thermo Fisher Scientific).EBs were grown in suspension culture for 8 days, then plated onto gelatin-coated plates and allowed to differentiate for an additional 8 days in DMEM + 10% FCS.

### Karyotyping

Chromosomal G-band analyses were performed at Nihon Gene Research Laboratories Inc.

### Image Acquisition and CNN analysis

Phase contrast and fluorescence images labeled with rBC2LCN-FITC (Wako, Osaka, Japan) were taken by the automatic laser-processing device (Kataoka Corp, Kyoto, Japan). 12,556 differentiated images and 18,834 undifferentiated images were used to train a CNN. The size of an input image was set to 70×70 pixels. Since the size of an original image is 1920×1200 pixels, local regions of 70×70 pixels can be cropped without overlap, and the regions can be fed into the CNN. Our CNN consisted of two convolutional layers, two pooling layers, and a fully connected layer. Thirty-two filters with 5×5 and 3×3 kernel sizes were used in the first and second convolutional layers, respectively. After the convolutional layers, we used a maximum pooling with 3×3 and 2×2 kernel sizes. The softmax cross entropy loss was used to train the CNN. Undifferentiated images were gathered in two ways. First, 70×70 local regions were cropped randomly from differentiated and undifferentiated regions, and we trained the CNN using those images. The trained CNN was applied to images, and misclassified undifferentiated regions were gathered. The gathered undifferentiated cell images were added to the training images, and the CNN was trained again and used as the classifier. The probability of obtaining a differentiated class from the CNN is assigned to the pixel in the resulting image.

### Automatic Cell Purification using LILACK system

hiPSCs (201B7 cell line) cultured in feeder-free conditions on light-responsive substrate-coated 35 mm dishes (Iwaki) for 7 days were used for laser irradiation. First, whole-dish tiled images were taken by the LILACK device (Kataoka Corporation) and were analysed by the algorithm made by the CNN analysis described above. Then, the laser was irradiated to the area automatically determined as differentiated cells by the algorithm. All the cells in the treated dish were used for flow cytometry analysis.

### Flow Cytometry

Harvested cells were fixed with PBS containing 4% (vol/vol) paraformaldehyde for 10 min at room temperature. Then, the cell samples were washed with PBS containing 0.5 mM EDTA and 1% (vol/vol) FBS and stained with mouse monoclonal anti-TRA1-60 antibodies, clone TRA1-60, Alexa Fluor 488 conjugate (Merck Millipore; MAB4360A4), for 1 h at 4 °C. Cells were then washed three times with PBS containing 0.5 mM EDTA and 1% FBS. After washing, cells were filtered through a 70 mm cell strainer (BD). The stained cells were analysed with FACSAria II (BD) and FACSDiva software (BD).

### Determining the Cell Numbers and Viability

The cell numbers and viability were examined using an automated cell counter (NucleoCounter NC-200, Chemometec). Briefly, total and dead cell numbers were counted by staining with acridine orange and DAPI, respectively.

### Viruses and Mycoplasma Infection Tests

Viruses and Mycoplasma Infection Tests, using multiplex quantitative PCR methods, were performed in PharmaBio Corporation.

## Acknowledgements

We thank Dr. Bruce R Conklin for scientific critical review This study was financially supported by a JSPS KAKENHI Grant-in-Aid for Scientific Research B (25282148, 16H03845) to K.S., a TIA (Tsukuba Innovation Arena) Kakehashi Grant to K.S. and Y.H., a Tsukuba Collaborative Project Grant to K.S. and Y.H., a JSPS KAKENHI Grant-in-Aid for Young Scientists (A) (17H05063) to Y.H., Grants for Regenerative Medicine, Japan Agency for Medical Research and Development (AMED) to Y.H., a Kowa Life Science Foundation Research Grant to Y.H., Takeda Science Foundation to Y.H., Uehara Memorial Foundation to Y.H., Tsukuba University Grant A and S to Y.H., Mochida Foundation to Y.H., Mother and Child Health Foundation to Y.H., and a Kyoto Sangyo 21 Research and Development Grant to Kataoka Corp, iPS Portal. Inc., and AIST.

## Competing Financial Interests

Y.H. was a former employee of iPS Portal, Inc. J.M. and M.S. are employees of Kataoka Corporation. L.X., Y.K., and S.H. are employees of iPS Portal, Inc. Y.H., K.H., and K.S. received a research grant from Kataoka Corporation.

## Author Contributions

K.S. and J.M. initiated the study. Y.H. developed the hiPSC assays and analysed the data with L.X., K.Y., and S.H. J.M. developed the laser irradiation and image acquisition and analysis system with M.S. K.H. developed the image processing and machine learning pipeline with S.K. K.S. developed the light-responsive polymer-mediated culture conditions with K.M. and T.K. Y.H., K.H., and K.S. supervised the study and wrote the manuscript. All authors commented on the manuscript.

